# Bioinorganic Angiogenesis

**DOI:** 10.1101/455212

**Authors:** Sophie Maillard, Baptiste Charbonnier, Omaer Sayed, Aslan Baradaran, Harshdeep Manga, Benjamin Dallison, Zishuai Zhang, Yu Ling Zhang, Sabah N.A. Hussain, Dominique Mayaki, Hermann Seitz, Edward J. Harvey, Mirko Gilardino, Uwe Gbureck, Nicholas Makhoul, Jake Barralet

## Abstract

Causing a large diameter blood vessel to sprout branches and a capillary network on demand to create a new angiosome is key to harnessing to potential of regenerative medicine and advancing reconstructive surgery. Currently this can only be achieved by connecting a vein graft to an artery by microsurgery, the arteriovenous loop technique (AVL). The arterial blood pressure in the thin-walled vein is thought to drive remodelling to create branches, however the surgical complexity limits the application of the technique. In this study we demonstrate that unexpectedly, a vessel density of luminal branches in excess of that achieved by the surgical AVL approach can be induced simply by placing a vein in contact with a microporous calcium phosphate. Only osteoinductive biomaterials have been reported previously, this is thought to be the first report of an angio-inductive material. Pilot studies indicated that the material type greatly affected the degree of luminal vascularization. Material contact with the vein is not a requirement for luminal angiogenesis of the vein and together these findings point to a bioinorganic effect, wherein the degradation of the material both releases a stimulatory ionic milieu and creates space for the developing angiosome.

## Introduction

Advances in regenerative medicine are hindered by problems caused by one central issue; the inability to direct and control blood vessels. The observation that materials alone, with no bioactives, cells or clinical pre-treatments can be invaded by microvessel capillary ingrowth is well documented. Capillary infiltration into pores as small as 1 µm is material independent,^[1]^ with explanatory hypotheses based mainly on inflammation and hypoxia-driven reaction of macrophages.^[2]^ Macrophages have been observed to migrate down oxygen concentrations releasing various growth factors (e.g., VEGF, IL-8, FGF1, FGF2)^[3,4]^ that can promote capillary invasion.^[5–7]^ However, the average capillary length is around a few hundred microns and a nutrient diffusion extends only a further 200 from a capillary ^[8,9]^ and so the depth of implanted materials that can be nourished by capillary ingrowth following implantation is limited. This means that only small or 2D sheet shaped tissue volumes can be sustained. A common example being split skin grafts.^[10,11]^ Too thick to be sustained by capillaries alone, the dermis is shaved to a faction of its normal thickness to allow tissue survival when transplanted.^[12]^

Branching of large (milli-scale) vessels can occur physiologically, for example in the development of collateral vessels in the case of atherosclerotic obstruction.^[13,14]^ Here the elevated pressure caused by the arterial narrowing is thought to drive remodelling of capillaries and minor vessels into collaterals. A surgical analogue of this process is created with the arteriovenous loop technique (AVL). Here, microsurgical anastomosis of a vein to an artery results in extensive luminal branching thought to be driven by arterial blood pressure (typically 65-110 mmHg) ^[15]^ causing remodelling in a thin walled vein that physiologically only carries blood at a pressure of 3-10 mmHg.^[16]^

Neovascularisation of a new angiosome, (i.e. a new blood vessel ‘tree’ fed by one large vessel that can sustain a defined tissue volume) is not considered possible in adults without using the AVL technique, but is the goal of many regenerative attempts to restore function, limit the consequences of ischemia and initiate enhanced healing. Milli-scale vessels are significant because the limit of a vessel that can be routinely transplanted using microscope assisted surgery is ~1mm.^[17]^

Given the overarching importance of being able to induce luminal sprouting from major blood vessels within a defined volume in order to sustain regenerating tissue volumes, surprisingly little has been published on interaction between biomaterials and major (milli-scale) blood vessels. Most work has focussed on blood-material interactions to develop non-thrombosing vessel substitutes.^[18–20]^ Attempts have been made to induce angiogenesis by delivering cytokines known to be involved in the process of blood vessel formation such as VEGF and FGF, endothelial cells and their precursors.^[21,22]^ While with some preclinical success these approaches are expensive and with undefined risk profile. Furthermore therapeutic angiogenesis has not as yet been found to be beneficial clinically.^[22]^ New blood vessels from using approaches to date do not last long and cannot oxygenate areas starved of oxygen. It has been shown that stimulation by non-angiogenic protein (Fibroblast growth factor 9) during angiogenesis created robust long lasting microvessels by stimulating the perivascular cells from the tunica adventitia and tunica media ^[23]^. Subsequently several studies have confirmed an important role for perivascular tissue in new vessel development.^[24–28]^

Given the striking importance of perivascular tissue and the lack of published interactions between the perivascular tissue and biomaterials we sought to investigate how a well characterised calcium phosphate bioceramic would interact with the tunica adventitia of a femoral vein. Here we report on the massive luminal sprouting that occurred when the tunica adventitia was encased by a microporous calcium phosphate material and compare this to a material simply implanted close to a vein and also a microsurgically created AVL.

## Results

### Implant characterization

Calcium phosphate scaffolds were successfully 3D-printed with an average deviation of + 150 µm when compared to their theoretical CAD model (**Figure S1**). Two circumferential grooves (200 µm thick, 300 µm deep) were carefully carved with a scalpel by hand prior to scaffold sterilization and implantation to retain sutures. Scaffolds were mainly composed of monetite (80.2%) and 19.8 % unreacted α- and β-TCP phases (**Figure 1A and B**). The porosity was bimodal about 2 and 5 micron modes (**Figure 1C and D**), and more than half the pore volume was in the range 1-10µm.

**Figure 1.**
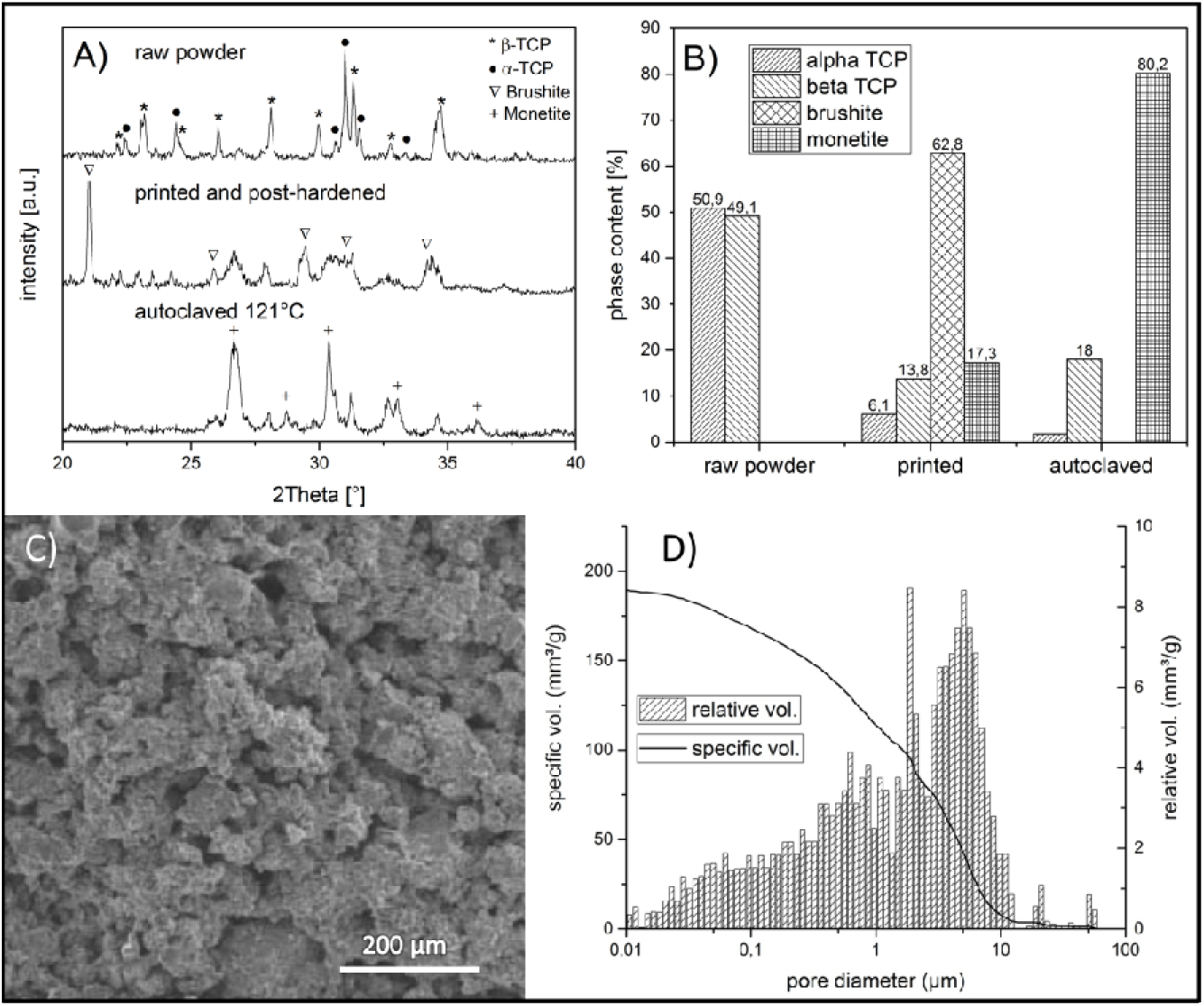
A) X-ray diffraction patterns of cement raw material, 3D printed samples and autoclaved implants. B) Quantification of phase composition by Rietveld refinement analysis, C) scanning electron micrograph of the surface of the printed monetite, D) histogram of pore size distribution and cumulative pore volume of printed monetite.

### Implant biodegradation and vascularization

The volume of the scaffolds decreased by ~15% after 4 weeks of implantation, whether or not they were pre-vascularized (**Figure 2**). While not significant (0.08 > p > 0.02), the mean scaffold biodegradation was greatest in the AVL group. The thickness of the implant walls decreased homogeneously throughout the implant (**Figure S1**) and the overall porosity also increased.

**Figure 2.**
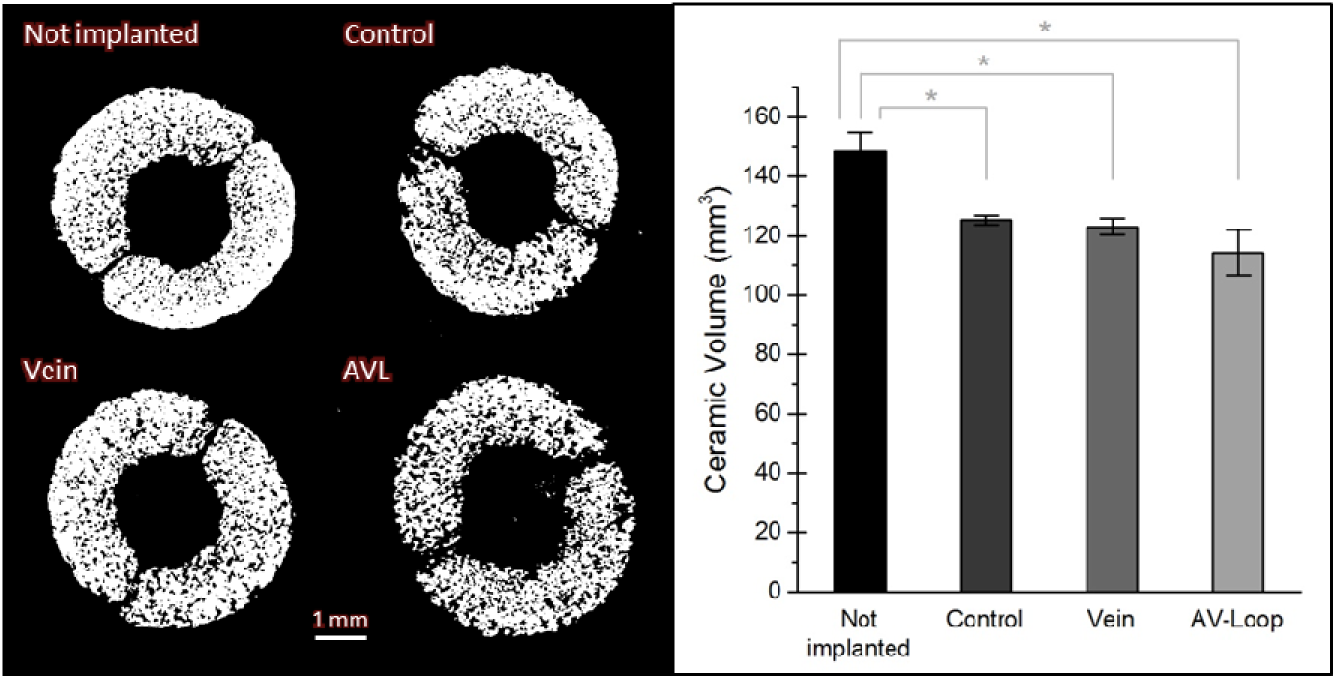
Degradation of the calcium phosphate scaffolds after 4 weeks of implantation evaluated through the analysis of µCT-scans of the explants. Illustrative cross-sections of the explants for each studied group are displayed on the left. (*corresponds to a statistical P <0.005)

Differences in morphology of vessels and invasion of the implants were observed between the different groups when examined by µCT of decalcified samples perfused with Microfil contrast agent and histology. A fibrous capsule and shell-like network composed of blood vessels was observed around the implant in all the groups (**Figure 3, 5 and S2**). Although the thickness of this capsule is sample-dependent and often variably distributed around the implant (e.g., **Figure 5 A1-C1**), the first layer of vascularization is often organized as follows: blood vessels between 300 to 500 µm in diameter grew along the bioceramics along their entire length. A secondary network of smaller vessels issued from their branching tended to colonize the surface of the implant along its periphery (e.g. **Figure 3C** longitudinal cross section). In the negative control group, vascularization of the implant originated from this vascularized capsule: the smaller vessels (150 µm < Ø) invaded the biocement porosity, sometimes reaching the middle of central channel (2 mm from the external surface). The presence of vessels with a diameter over 300 µm inside the central channel was observed occasionally, entering the implants from the edges, crossing them entirely and generating a network of smaller vessels. However, the density of blood vessels within the scaffolds remained quite low in comparison with the AVL and vein groups (**Figure 4, p > 0.005**). The small gap between the two halves of the ceramic scaffold appeared to be a preferential site for vascular invasion of the central channel of the scaffold (e.g. **Figure 3A**, transversal cross section). Even though the role played by this fibrous capsule cannot be disregarded, the presence of an axial vascularization resulted in a dramatic increase (~ x 1.6) of blood vessel density inside the scaffold (~ x 2.0 for vein) (**Figure 4, p > 0.005**). For the AVL group, a splitting of the main axial vascularization (Ø ≈ 600 - 1200 µm) was often observed (**Figure 3B**), branching into several lower diameter axial vessels (Ø ≈ 400 - 600 µm). A secondary network of vessels (Ø ≈ 150 - 350 µm), sprouting from theses main vessels axially colonized the scaffold’s central channel, allowing for the invasion of the bioceramic by smaller vessels (25 < Ø < 150 µm) and capillaries (25 µm < Ø). The morphology of the vasculature generated within the scaffolds with central veins was slightly different. The vein itself did not branch, and numerous venules (Ø ≈ 150 - 400 µm), surrounded it (**Figure 3C and 5**). This radial vascularization of the implant by smaller vessels issuing from these venules, formed a high density vascular network. In general, for both AVL and vein groups, no large diameter vessels (Ø > 300 µm) were observed invading directly the porous structure of the bioceramic, however, some could be found in the volume where the scaffold structure had degraded

**Figure 3.**
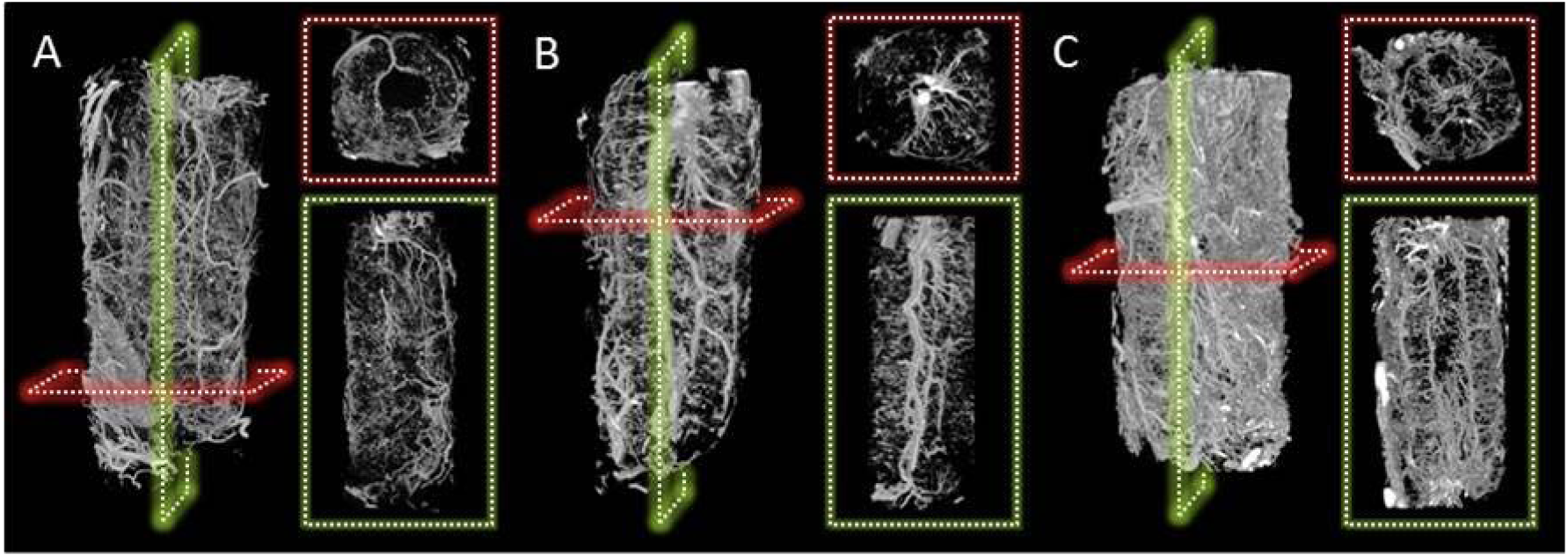
Vascular networks developed within and next to the calcium phosphate scaffold after 4 weeks of implantation, as shown by micro-CT of the decalcified explants, A) negative control group (without axial vascularization), B) AVL group and C) vein group. Example replicate images **Figure S2**

**Figure 4.**
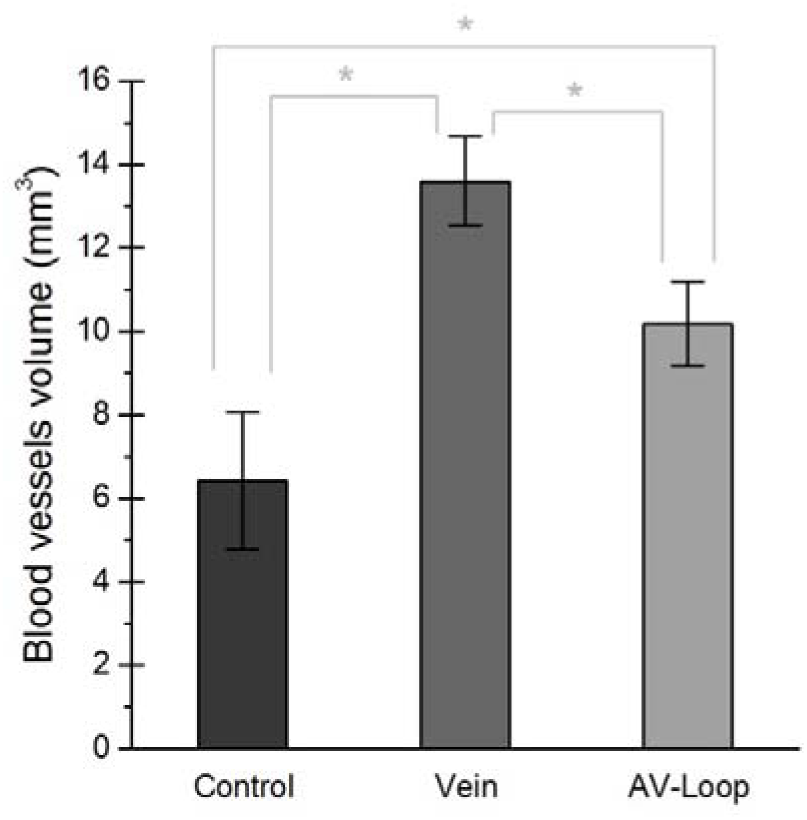
Quantification of blood vessel volume inside the calcium phosphate scaffolds after 4 weeks of implantation determined through micro-CT-scan analyses of decalcified explants. Special care was taken as to the definition of the region of interest to consider avoiding inclusion of the fibrous capsule. (* P <0.005)

**Figure 5.**
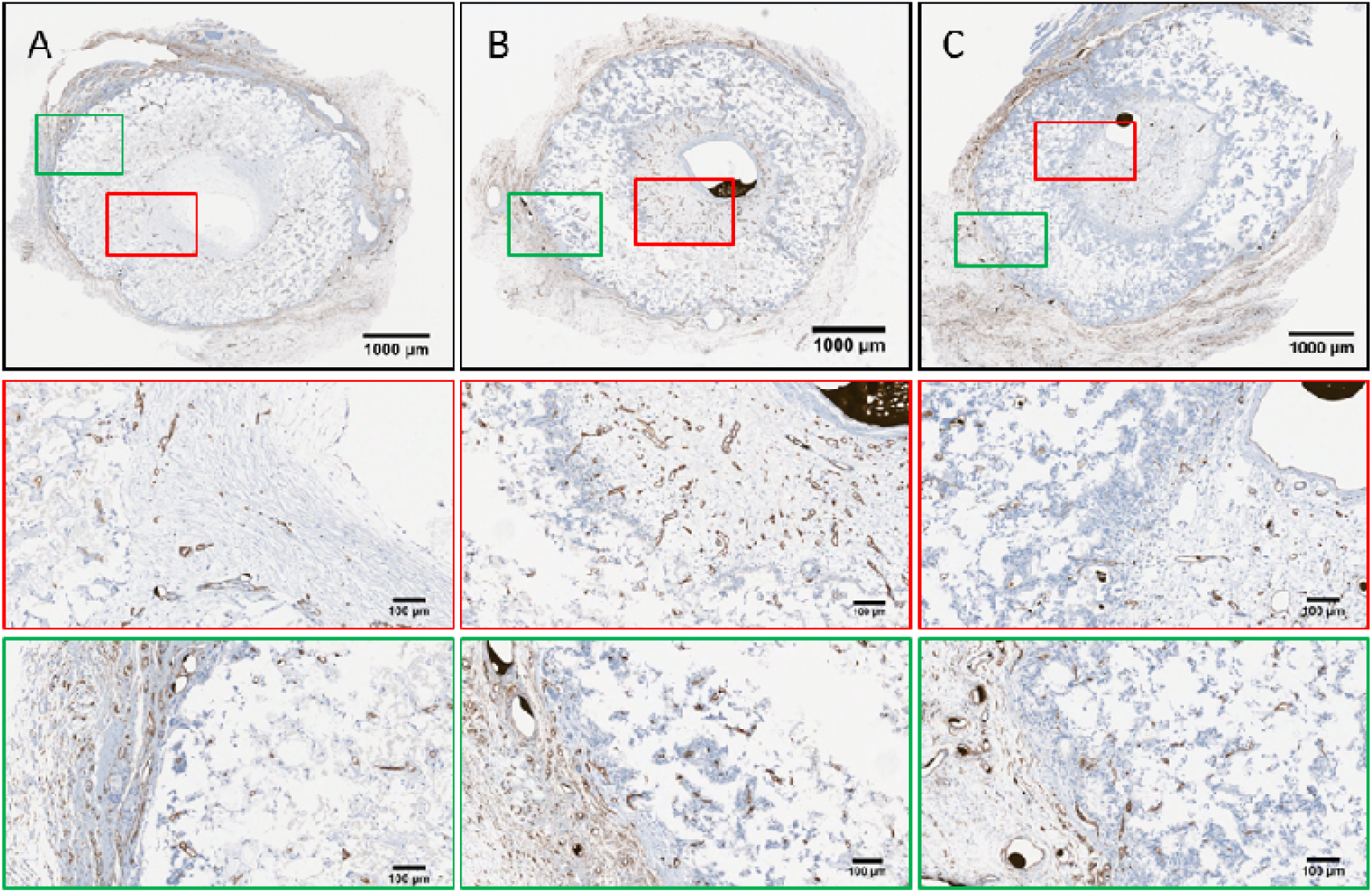
CD34 immunohistological sectoins of A) negative control group, B) vein group and C) arteriovenous groups. A similar illustration is available (**Figure S3**) showing the same regions but with an H&E staining.

Histological observations illustrated in **Figure 5 and S3** confirmed µCT measurements. In all groups the highly vascularized capsule was present, at the surface of which muscle tissue could be observed (**Figure S3A and C**). Arterioles or venules were present in this capsule. H&E staining (**Figure S3**) revealed that the organization of the vessels inside the scaffold channel was determined by whether or not the scaffold had axial perfusion. In the control group, the tissue, which filled only partially the channel, was poorly vascularized and composed of collagen layers that were deposited parallel to the implant’s internal surface. The density of blood vessels was much higher in the case of the AVL or vein axial perfusion, and the channel was full of fibrous tissue. CD34 staining revealed that the vessels sprouting from the central vein possessed the characteristics of venules, i.e. thin outer wall and a large lumen (**Figure 5B and C**). A higher number of blood vessels in the vein group compared to the AVL loop and control groups was observed in in CD34 stained histological slices – and similarly between the AVL and control groups.

No TRAP activity was detected inside or outside the bioceramics (**Figure S4**), regardless of the experimental group, which indicated the osteoclasts were not responsible for the resorption of the calcium phosphate implants. The revascularization of the scaffolds axially perfused by a vein and isolated from the surrounding environment by a silicone sheet was greatly delayed if not impaired. Indeed, blood vessel density after 4 weeks of implantation was 2 to 3 times lower than for the vein group (p < 0.005), and even below the value determined for the non-perfused control group (**Figure 7 and S5B**).

### Gene expression – axial vein model

The expression of genes known to play a role in the formation of blood vessels was determined and compared between a native vein, a distal segment of the same vein inside the bioceramic tube with and without a silicone sheet wrapped around. RNA was extracted from within the bioceramic. RT-qPCR gene expression is shown in **Figure 6**. It is notable that only gene expression inside the scaffolds was higher than vein tissues (p < 0.005).

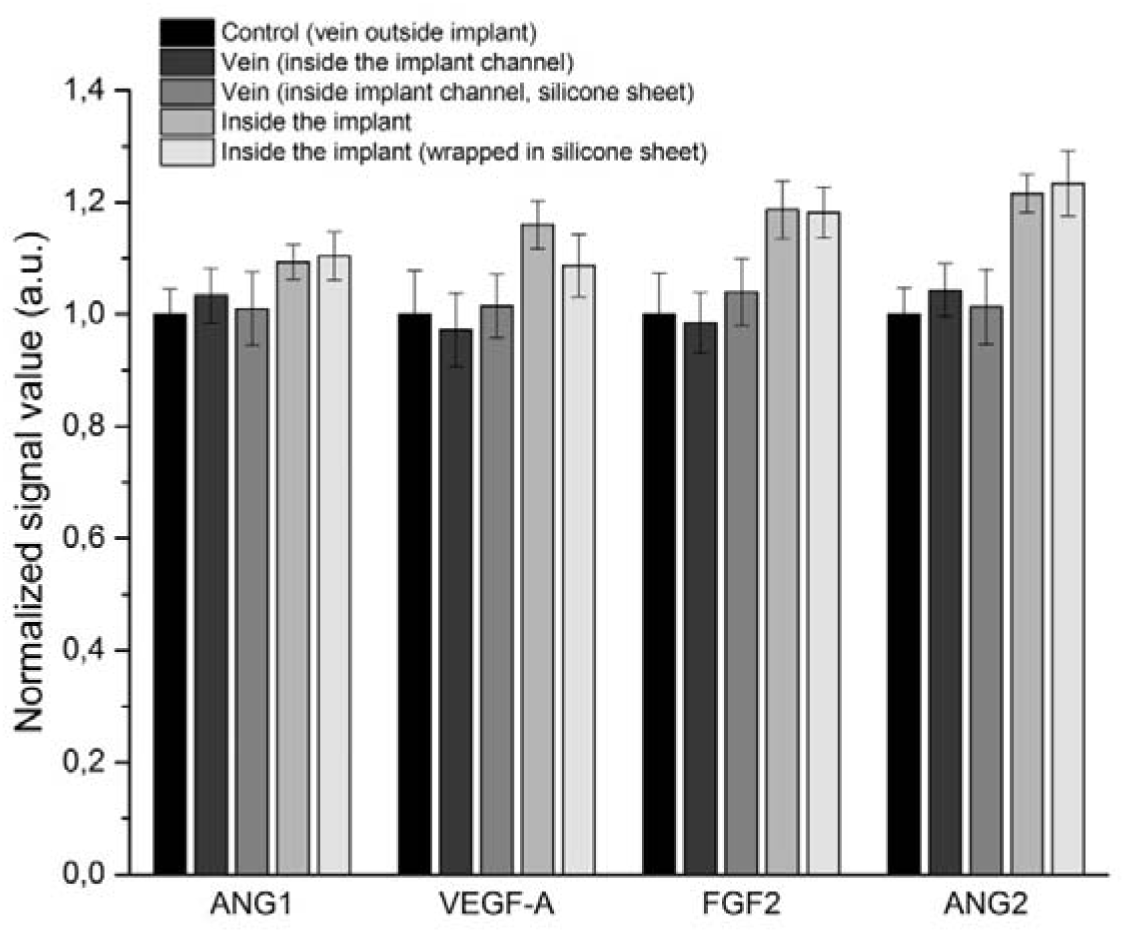
Figure 6. Gene expression of Angiopoietine 1 (ANG1), vascular endothelial growth factor A (VEGF-A), fibroblast growth factor 2 (FGF2) and angiopoietin 2 (ANG2) determined by quantitative real time. reverse polymerase chain reaction in different location of implants perfused with a vein and wrapped or not in a silicone sheet. The signals were normalized according to the expression of the cyclophilin.

## Discussion

Within the biomaterials literature the pro-angiogenic effect of microporous polymers towards microcapillaries was widely investigated in the 1980s for grafts and tissue guidance membranes.^[2,29,30]^ Mechanisms driving the vascularization of microporous structures are not well understood and even for a single material, vascularization can depend on host-biomaterial interactions, stabilization of the scaffold after implantation and its local surrounding environment.^[2,31]^

We observed that circumferential contact with microporous ceramic induced luminal sprouting and so speculated that like capillary invasion of microporous polymers, the effect should be material independent. We posited that if microporosity of the calcium phosphate was driving the lumen of the blood vessels to branch, then analogous polymer structures would also. A pilot study comprising polymeric scaffolds (3D printed calcium phosphate, 75% microporous polycaprolactone (PCL) made by salt porogen (sieved to >75µm) leaching and 3D printed polymethyl methacrylate ^[32]^) was performed in the presence of a vein placed in the implant channel. After 4 weeks, although some vascularization was measured it was greatly reduced compared with monetite,(**Figure 7 and S6**) with a blood vessel density ~1.5 times lower than for the bioceramic control without any axial vascularization (p < 0.005).

**Figure 7.**
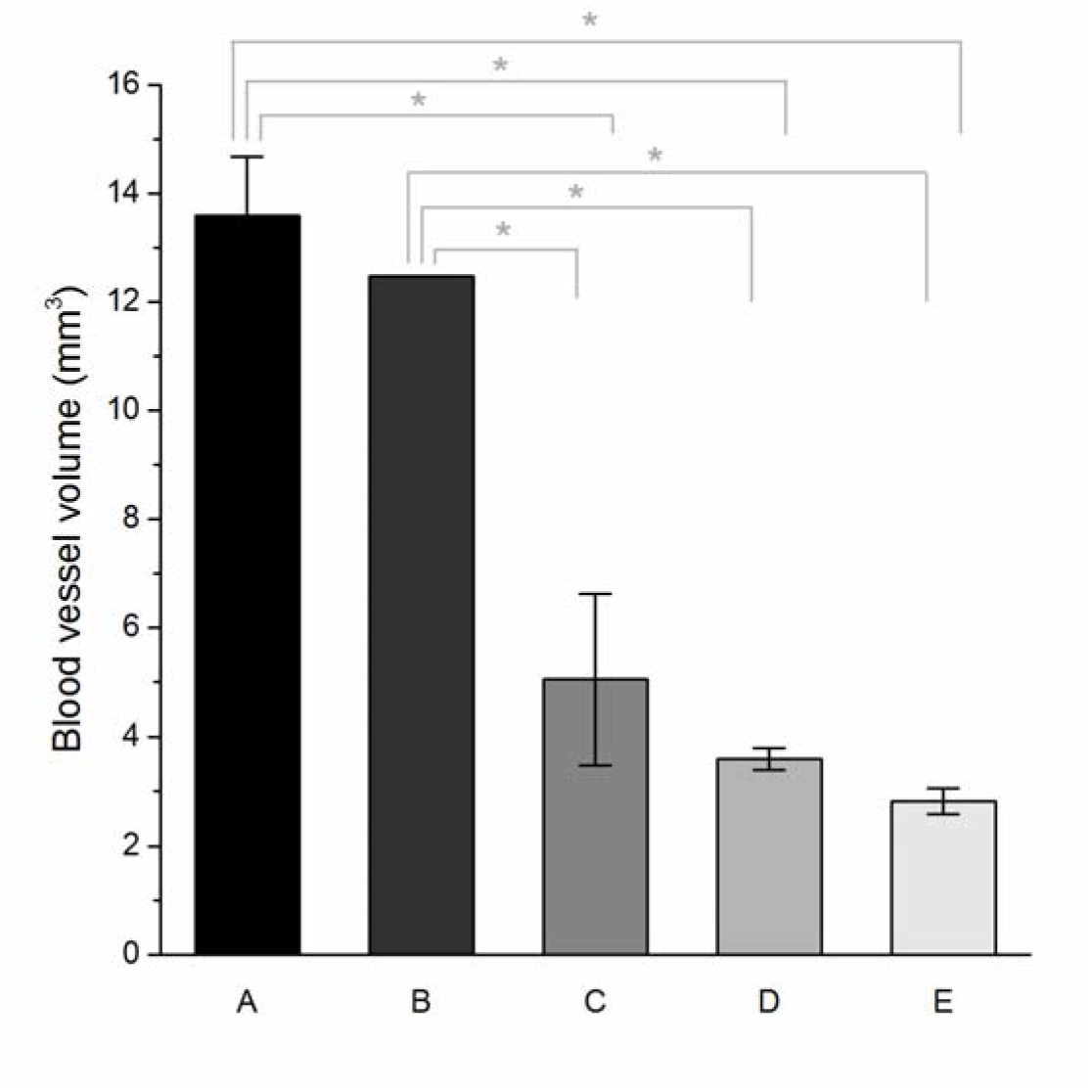
Comparison of the blood vessel density within implants of different compositions after 4 weeks of implantation. All the implants comported a venous axial vascularization. The 3D printed bioceramics (A) serving in this case as positive control. 3D printed bioceramics (A) were carved by hand in order to generate some internal groves (patterned bioceramic, B). The polymeric scaffolds D and E were respectively made of polycaprolactone (PCL) and polymethyl methacrylate (PMMA). (* P <0.005)

Relative hydrophobicity of the polymers may simply have reduced cell attachment and ingrowth since luminal sprouting was not eliminated in the polymeric scaffolds.

Inflammation has been suggested as another mediator of angiogenesis. ^[5,33]^ We sought to establish whether direct contact with the vessel was required for luminal sprouting. Inside monetite implants we carved away the internal surface of the implants creating six 2.5 mm square pegs close to the central vessel, leaving hollow chambers around the central vessel. (**Figure S5A**). Micro-CT indicated that at the millimeter scale, contact was not required for luminal sprouting. This finding, combined with the reduced angiogenesis in much less soluble polymer implants pointed to a role of a chemical effect.

Indeed a pro-angiogenic effect of calcium ions has been reported ^[34,35]^ While not conclusive, these observations are suggestive of chemical effect. ^[36–39]^ The ability of the monetite to resorb, apparently due to dissolution (**Figure S4**), may have also simply allowed more and larger blood vessel ingrowth. Indeed, the degradation of a scaffold at a rate matching the development of a new tissue is often considered as an ideal requirement.^[40]^ However, degradation rate alone is insufficient, for example if degradation products affect pH, this may inhibit angiogenesis.^[41,42]^Osteoclasts are blood derived cells and it was feasible that they could actively remodel the scaffold. TRAP staining was negative and so we infer that the increased resorption of vascularised cement occurred as a result of higher fluid turnover rates affected by the higher number of capillary networks.

The presence of a physical barrier limiting capillary ingrowth into the ceramics towards the lumen led to a decrease of blood vessel density by less than half that of the unwrapped implants (**Figure 7C**) indicating that some signalling from peripherally invading capillaries may have driven luminal sprouting. Unexpectedly, no significant differences in angiogenic gene expression were found in scaffolds with or without a silicone membrane (**Figure 6**).This may simply have been that vasculature inside the channel was mature at 4 weeks and that the formation of a new vascular network (e.g., ANG2) and its remodeling or maintenance (e.g., ANG1) occurred within the bioceramic walls.^[43,44]^

## Conclusion

This study reports three important findings. Firstly, the degree of neovascularization inside microporous calcium phosphate scaffolds is higher with a central axial vein than when placed around an arterially anastomosed AVL. This is highly significant as it indicates a non-microsurgical approach to vascularising regenerated tissue volumes. Secondly the material type strongly affects this luminal sprouting of the vein but microscale geometry does not. This is unexpected given there are reports in the literature that indicate different polymer types are similar at inducing capillary ingrowth, but there have been no ceramic polymer comparisons. Finally, it appears that extrinsic capillary ingrowth is a requirement for luminal sprouting of veins, possibly to provide the arterial half of the capillary bed.

## Experimental section

### Scaffold design and manufacture

The implants were designed using Alibre design Xpress 10.0 CAD software. As illustrated **Figure 8**, these cylindrical implants (12 mm long, 4 mm diameter) comprised a central channel of 2 mm diameter where a vessel could fit unrestricted but in close contact circumferentially (Figure 1a). Accordingly, to enable placement, the implants were designed in two halves which could be closed with sutures after the introduction of the blood vessel.

**Figure 8.**
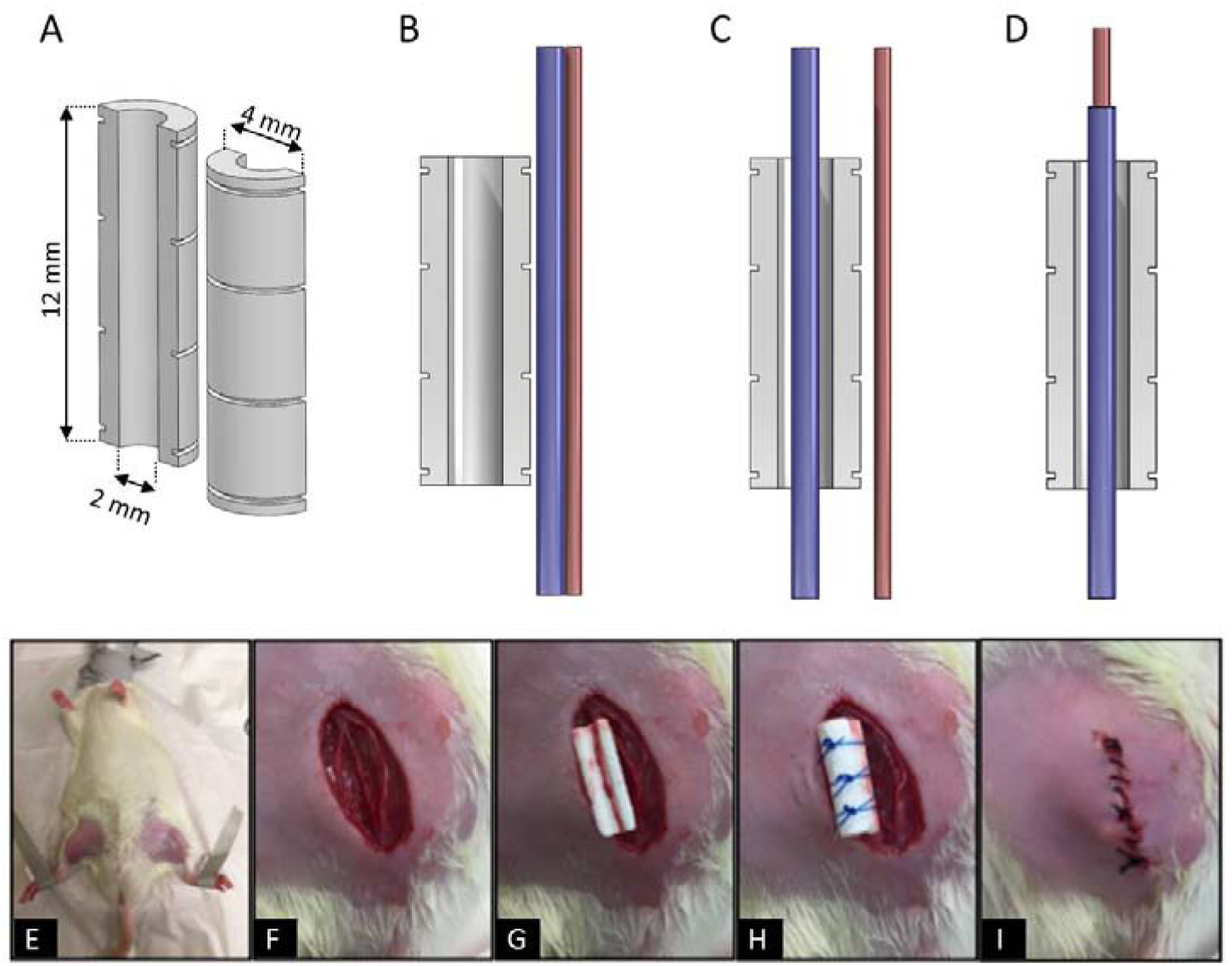
A) Computer-aided design of the printed scaffold and B) to D) schematic diagrams depicting the groups investigated: control, vein, arterio-venous loop, respectively. Photographs of the surgery show preparation of the surgical sites E), dissection of the femoral vascular bundle F), placement of the scaffold G), suturing the two implant halves around the vein H), and the closure of the incision I).

**Table 2.**
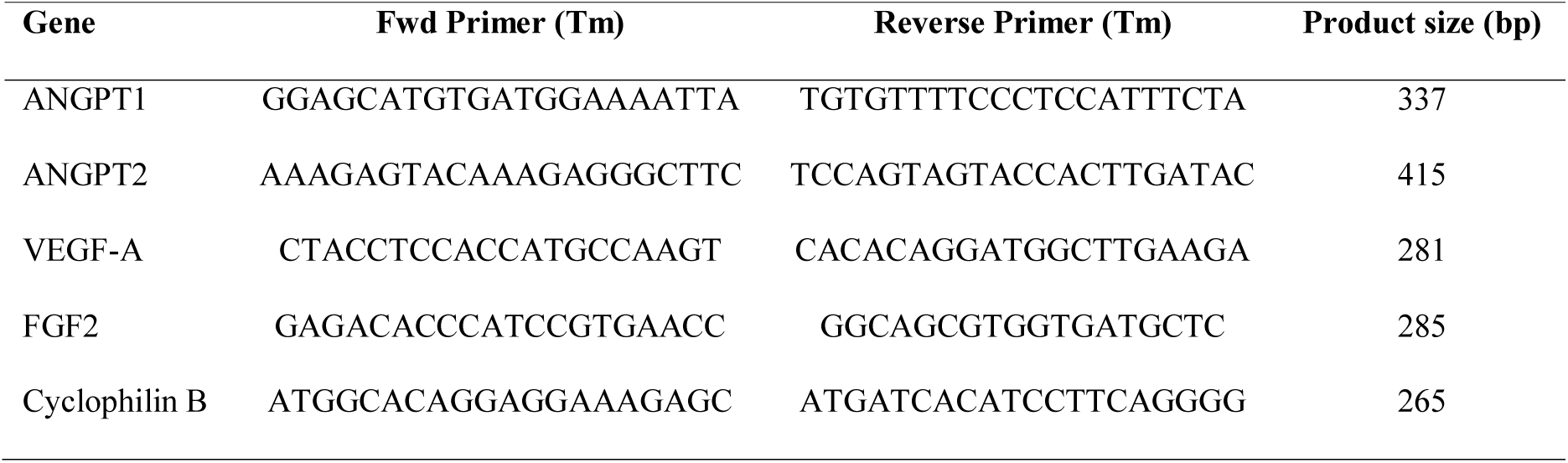
Primers used for Real-Time PCR experiments

Calcium phosphate scaffolds were produced by additive manufacturing according to a reactive 3D-printing technique co-developed by the authors.^[45,46]^ Briefly, a reaction between tricalcium phosphate powder (TCP, *Ca_3_(PO_4_)_2_*) and diluted phosphoric acid (*H_3_PO_4_*) allow for the area selective binding of the powder grains. After printing, samples are soaked in 20% phosphoric acid for 60 s, washed and sterilised by autoclaving.^[45–47]^

### Scaffold characterization

X-ray diffraction patterns of cement raw materials and printed implants were recorded with a Siemens D5005 diffractometer (Siemens, Karlsruhe, Germany) in a 2θ range of 20-40 °, a step size of 0.02 ° and a total measuring time of 3 s/step. The diffraction patterns were quantified by Rietveld refinement analysis using the software TOPAS 2.0 (Bruker AXS, Karlsruhe, Germany) with the internal database structures of alpha-TCP, beta-TCP, brushite and monetite as references.

Scaffold architecture was investigated by micro-tomography X (SkyScan 1172; SkyScan Kontich, Belgium) at a resolution of 12 μm and with a 0.5 mm aluminum filter. After reconstruction with NRecon (Skyscan), the difference in dimensions between the printed scaffold and the CAD model were evaluated using CloudCompare (EDF R&D). The microstructure of the implant was studied using scanning electron microscopy (SEM, Hitachi S-4700 FE-SEM; Tokyo, Japan) in back-scattered electron (BSE) mode at an accelerating voltage of 20 kV to enable visualization and quantification of the scaffold. The porosity and pore-size distribution of the 3D printed implants was determined by Hg porosimetry (PASCAL 140/440, Porotec GmbH, Hofheim, Germany).

### Animal model and surgical procedure

The animal study was performed with a total of 25 male Wistar rats (200-300 g, 49-56 days) after approval from McGill University Animal Care Committee (UACC). Femoral sites were considered for surgery because of their ease of access and presence of the femoral vascular bundle. Briefly, after anesthesia and analgesia, a skin medial incision allowed to expose the femoral vessels between the inguinal ligaments proximally and the bifurcation of the saphenous and popliteal vessels distally. For the positive control group (AVL group, N = 5) blood vessels were dissected. The tunica adventitia and media of the artery were removed without injuring the tunica interna. The femoral artery was anastomosed with the femoral vein to create the AVL. The first half on the scaffold was positioned, the AVL was positioned in the channel and the second half of the scaffold was placed and sutured to form a cylinder, with a special care not to pinch the AVL. A similar procedure was performed for the vein group (N = 6) where the femoral vein was inserted inside the bioceramic, respectively. A group wherein a scaffold was placed adjacent to a vein served as negative control (N = 4). **Figure 8** illustrates the three experimental groups (B to D) as well as the surgical procedure (E to I). Following the implantation, subcutaneous and cutaneous ligations were performed with Monofilament 4.0 to close the wound. Saline was administered depending on the volume of blood loss.

After 4 weeks of implantation, animals were perfused by a contrast agent (Microfil). In brief, perfusion was performed under anesthesia by cannulating the left ventricle with 150 mL heparin solution and 4% paraformaldehyde solution, leading to their death. The abdominal aorta was perfused with 10 mL of yellow Microfil (MV-122) containing 4% of MV Curing Agent (Flowtech). Afterwards, the abdominal aorta and cava vein were ligated and the animals were placed at 4°C overnight. The implants were finally explanted and the tissues were fixed in in 4% paraformaldehyde solution and animals were perfused by a contrast agent (Microfil). In brief, the later were perfused under anesthesia by cannulating the left ventricle with 150 mL heparin solution and 4% paraformaldehyde solution, leading to their death. The abdominal aorta was perfused with 10 mL of yellow Microfil (MV-122) containing 4% of MV Curing Agent (Flowtech). Afterwards, the abdominal aorta and cava vein were ligated and the animals were placed at 4°C overnight. The implants were finally explanted and the tissues were fixed in in 4% paraformaldehyde solution and µCT measurement allowed analysis of implant volume. Decalcification was performed in ethylenediaminetetraacetic acid (EDTA, 14%_wt_) at pH 7.2 for three weeks at 4°C until the samples were radiolucent at which point µCT was performed. Dehydration in ascending serial ethanol solutions preceded paraffin embedding, followed by sectioning into 5µm histological slices with a microtome (RM 2255 microtome Leica Microsystems, Germany), followed by haematoxylin and eosin (H&E) staining, CD34 for blood vessels (Abcam ab81289), and Tartrate resistant alkaline phosphatase (Sigma-Aldrich 387A-1KT). To investigate the effect of the surrounding environment on vascularization, ten additional rats underwent surgery according to the single vein model described previously; half of them having their implant wrapped in a silicone sheet before suturing the wound. After sacrifice (4 weeks of implantation), µCT analyses were performed. The underlying mechanisms driving scaffolds revascularization when axially perfused by a vein were also examined by quantitative real time polymeric chain reaction (Qrt-PRC, described in detail in a following section).

### Analysis of the explants

Implant biodegradation and vascularization were quantified using CT-Analyzer software (Bruker) on calcified and decalcified samples, respectively. Histological analysis was performed using a Ziess microscope Axio Imager.M2 (Zeiss Gottingen, Germany) with a digital AxioCam IC camera (Zeiss Gottingen, Germany).

### Gene expression – single vein model

Total RNA was isolated from the samples using TRIzol reagent (Cat. No. 15596-018; Invitrogen™, Carlsbad, CA, USA) following the manufacturer’s protocol and quantified in a spectrophotometer by absorbance readings at 260 nm. Three interest zones were targeted, i.e. in the bioceramic lumen where the vein was inserted and inside and outside the bioceramic, respectively; the control being an aliquot of the vein sampled outside the scaffold. 1 µg of total RNA from each sample was reverse transcribed using a high-capacity cDNA reverse transcription kit (Cat. No. 74322171; Applied Biosystems, Foster City, CA, USA) in accordance with the manufacturer’s instructions. The resulting cDNAs were used for real-time PCR using Power SYBR Green PCR Master Mix (Cat. No. 4367659; Applied Biosystems). Reactions were carried out in a 7500 real-time PCR system (Applied Biosystems) for 40 cycles (95°C for 15 s, 60°C for 30 s and 72°C for 45 s) after the initial 10 min incubation at 95°C. A Ct (cycle threshold) value for each reaction was calculated using Applied Biosystems sequence detections software and the relative ratio of expression was determined using a previously described algorithm.^[48]^ Primers used to amplify specific targets are as follows: Angiopoietin-1 (ANGPT1, Ang-1), vascular endothelial growth factor (VEGF-A), FGF2 (fibroblast growth factor 2) and angiopoietin-2 (ANGPT2, Ang-2) (Table 1). All mRNA expression values were normalized to the results obtained for cyclophilin mRNA (housekeeping gene).^[49]^

### Statistical analysis

Data are presented as representative images, representative experiments, or as means ± standard error of the mean, with N indicating the number of independent experiments. Statistical analysis was performed using StatPage calculator (http://statpages.info/anova1sm.html) with one-way analysis of variance ANOVA (turkey post hoc test), and a P <0.005.

## Acknowledgments

La Fondation des Gueules Cassées for provision of studentship (SM), NSERC CREATE Award (BC, OS), Quebec Government Bavaria-Quebec collaboration award, as well as the Fonds de Recherche du Nature et Technologies, (JB), NSERC discovery award (JB) and infrastructure support of JR Hayward grant, Chamber Lyons Academy.

Received: ((will be filled in by the editorial staff))

Revised: ((will be filled in by the editorial staff))

Published online: ((will be filled in by the editorial staff))

## Supporting Information

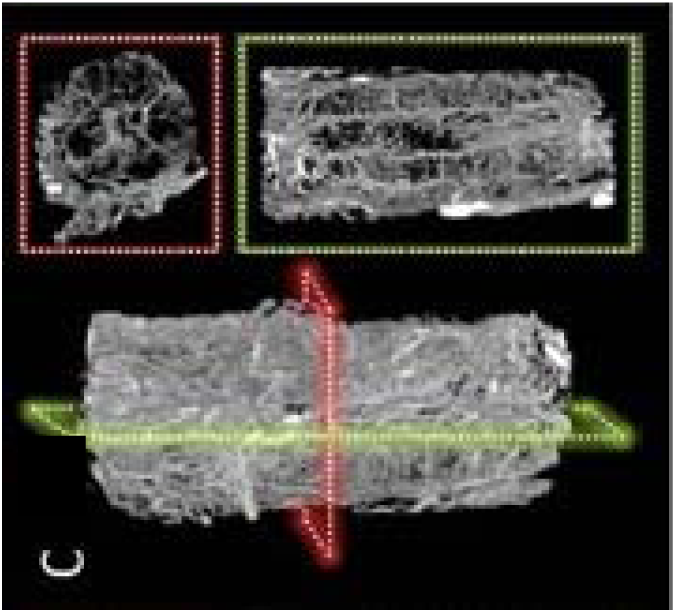

**Figure S1.**
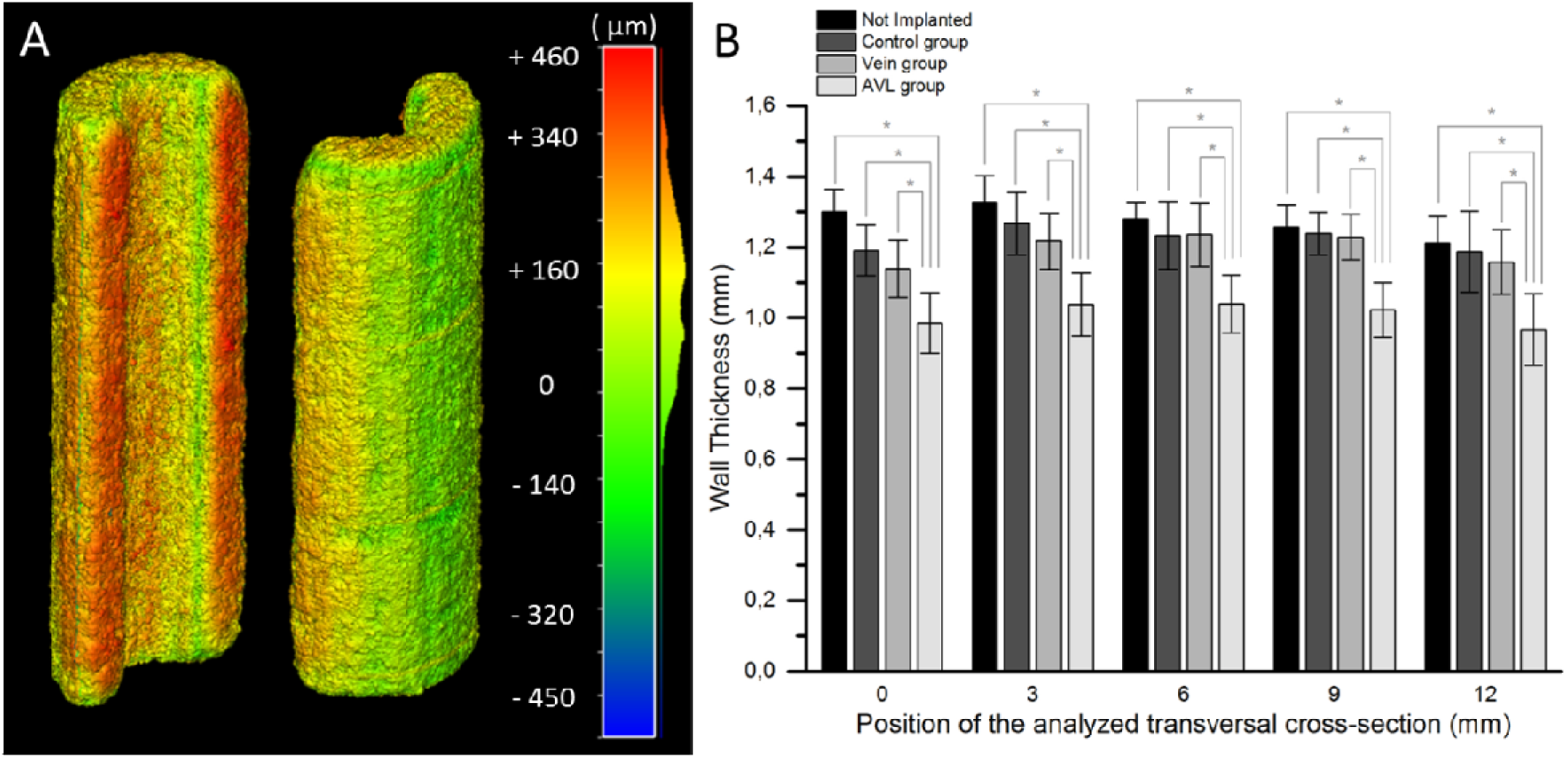
A) Coloured illustration of the geometrical deviation between the CAD design and the 3D printing ceramic (average + 150 µm, Gaussian fitting) and B) Evolution of the scaffold wall thickness determined by analysis of micro-CT transversal cross sections at different locations

**Figure S2.**
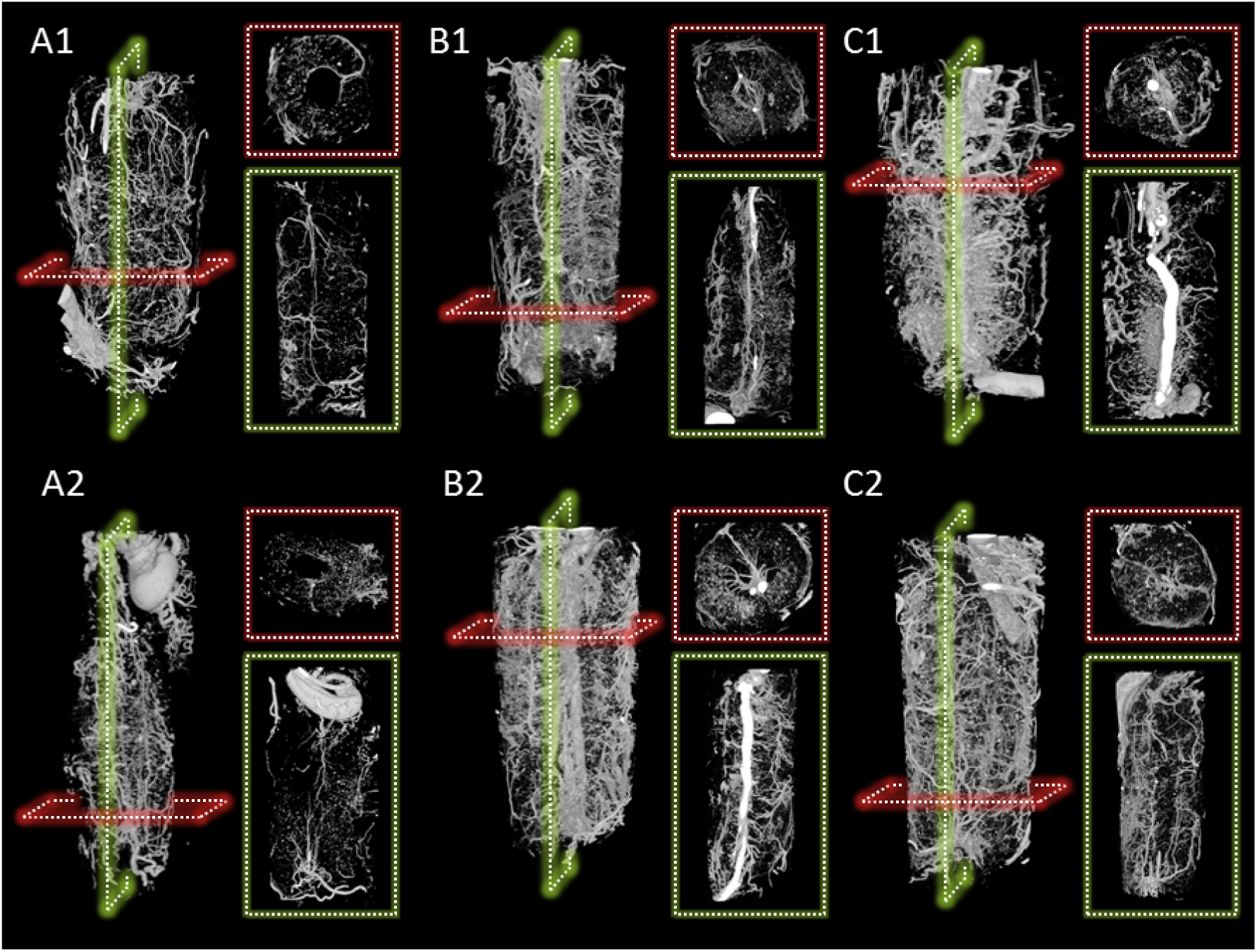
Replicate samples showing vascular network developed within and next to the calcium phosphate scaffold after 4 weeks of implantation, obtained by micro-CT of the decalcified explants, with A) the control group (without axial vascularization), B) the AVL group and C) the vein.

**Figure S3.**
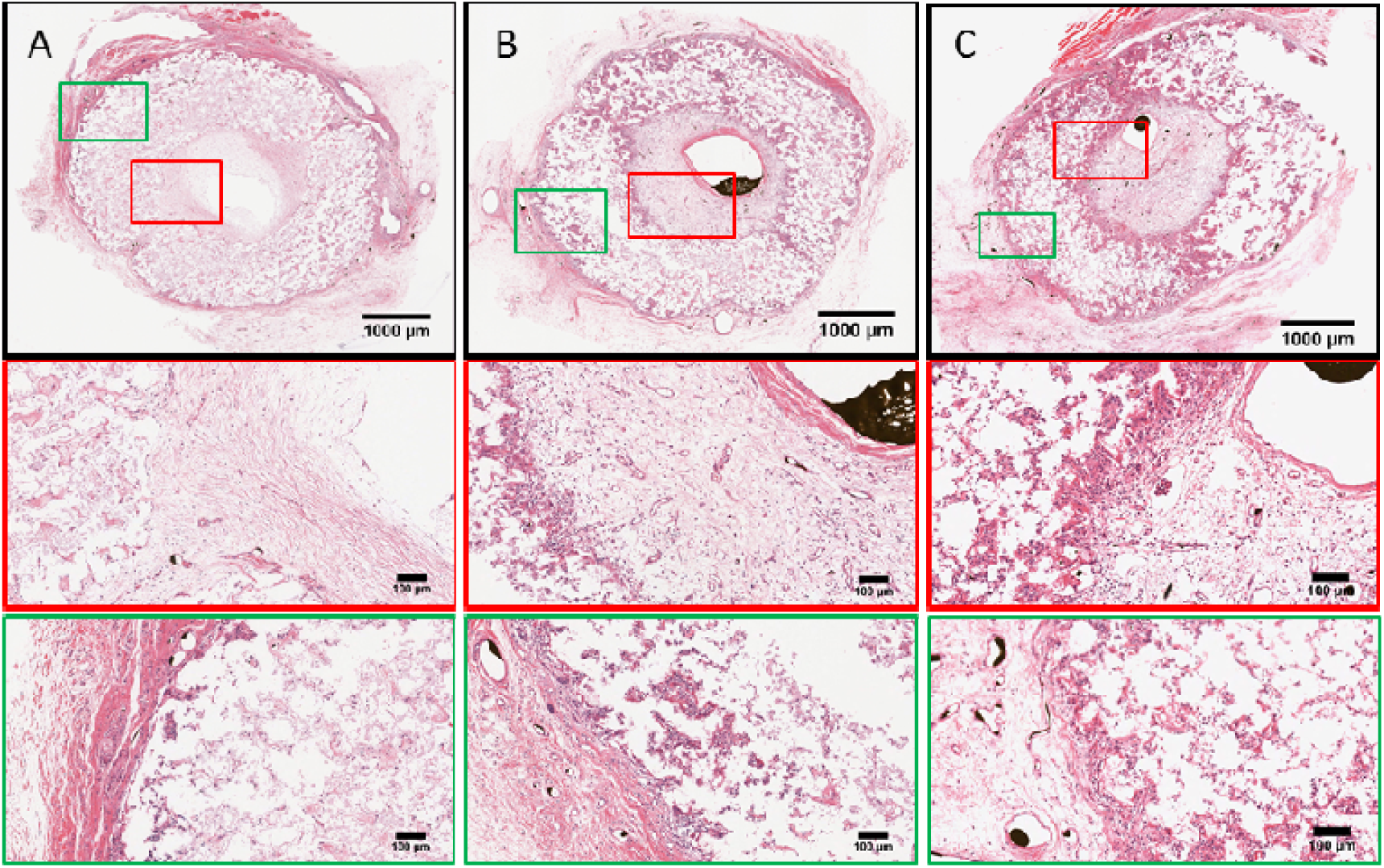
Optical microscopy images of H&E stained histology slices of A. the control group, B. the vein group and C. the AVL group. Enhanced magnifications of the green and red zones are also displayed, focusing on the inside and outside of the implant, respectively.

**Figure S4.**
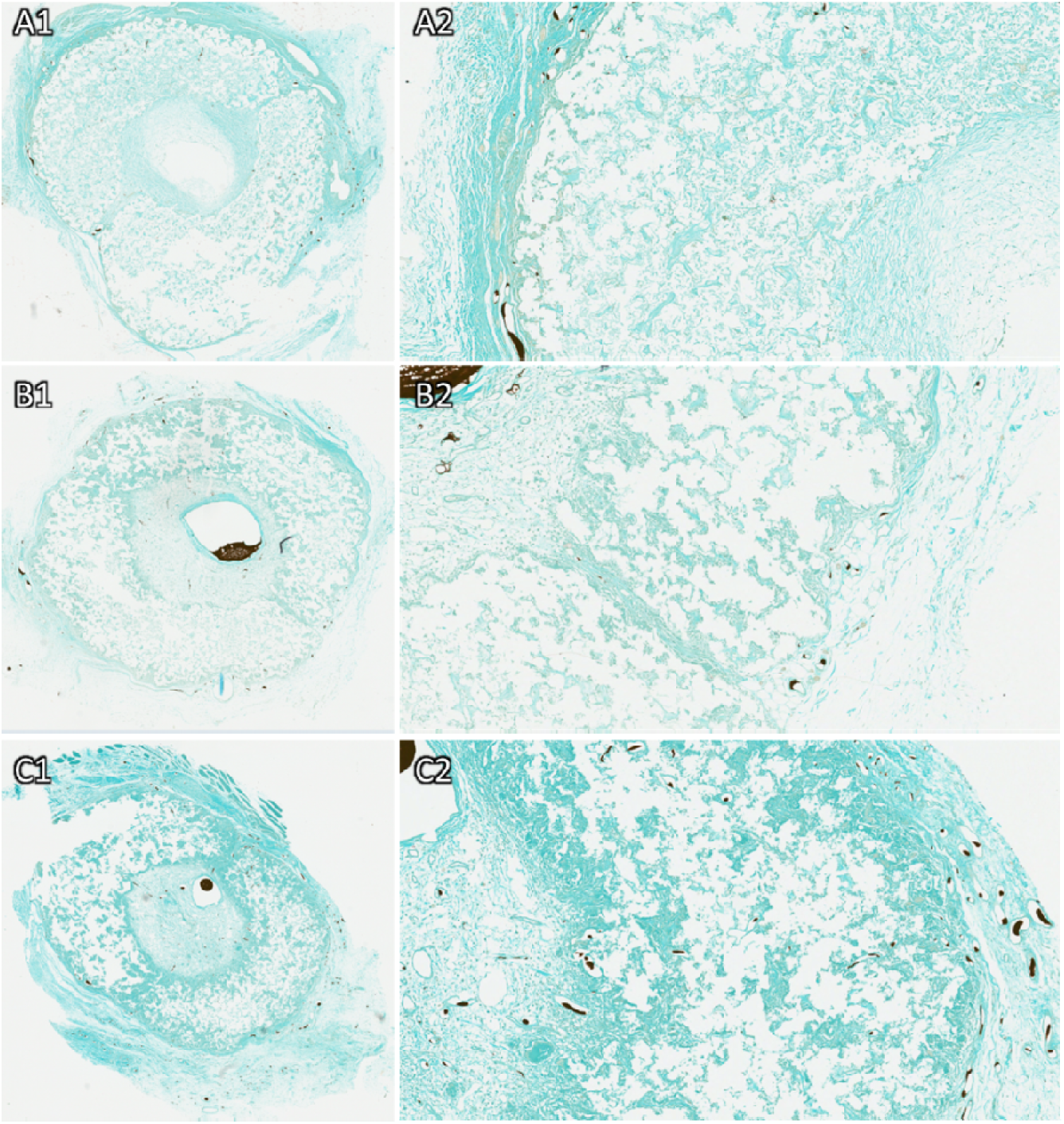
Negative TRAP staining of histological sections of A) control group, B) the vein group and C) AVL group. No osteoclast activity was observed (red color) in any experimental implants.

**Figure S5.**
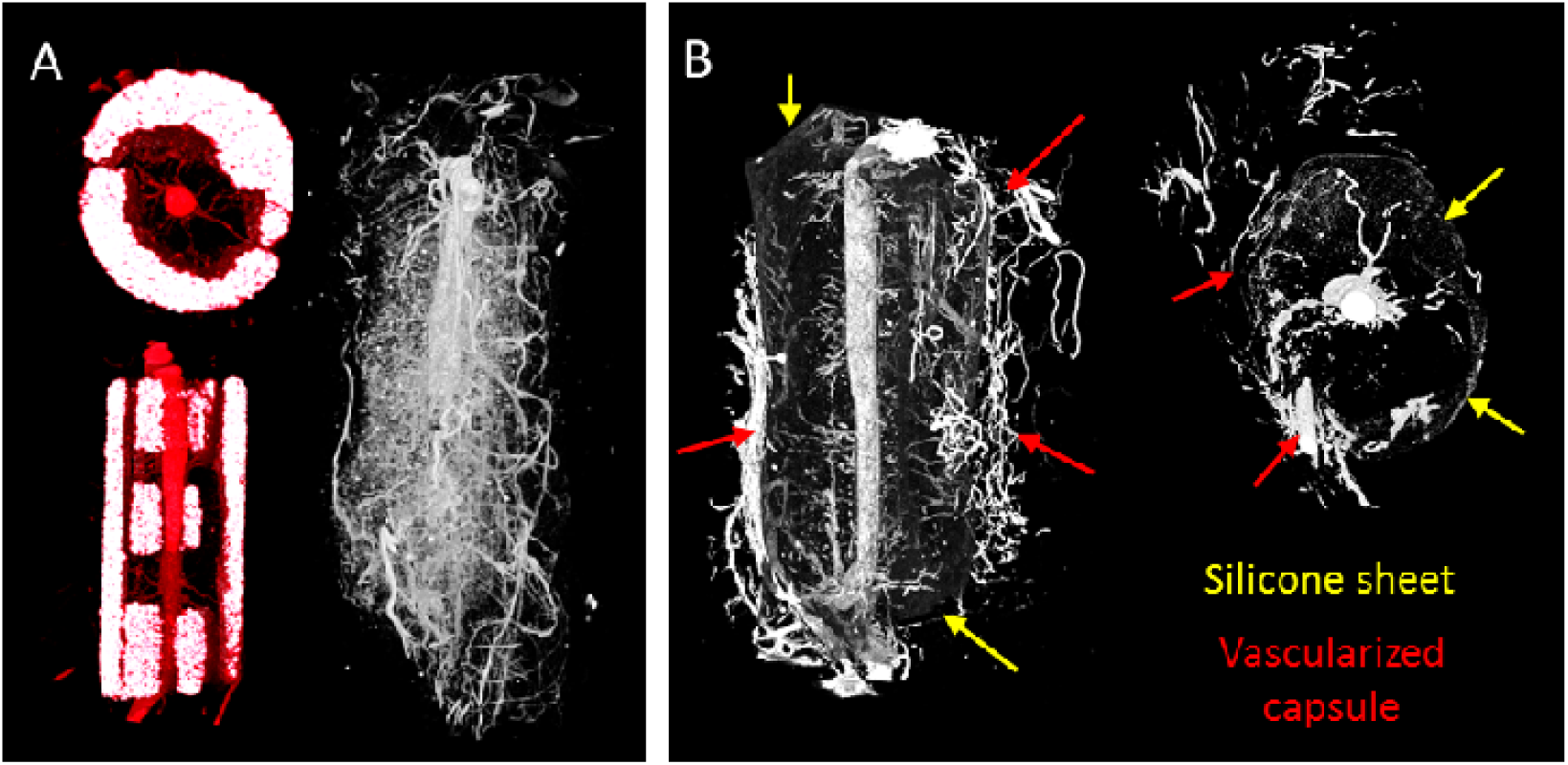
Micro-CT images of the vascularization inside carved calcium phosphate scaffolds before (left) and after (right) decalcification (A). No correlation with macroscale geometry and luminal sprouting was observed. (B) 3D µCT image illustrating paucity of luminal sprouting when extrinsic capillary ingrowth blocked by silicone membrane. A well vascularised fibrous capsule was formed (red arrows) that did not progress beyond the silicone sheet, (yellow arrows mark approximate position).

**Figure S6.**
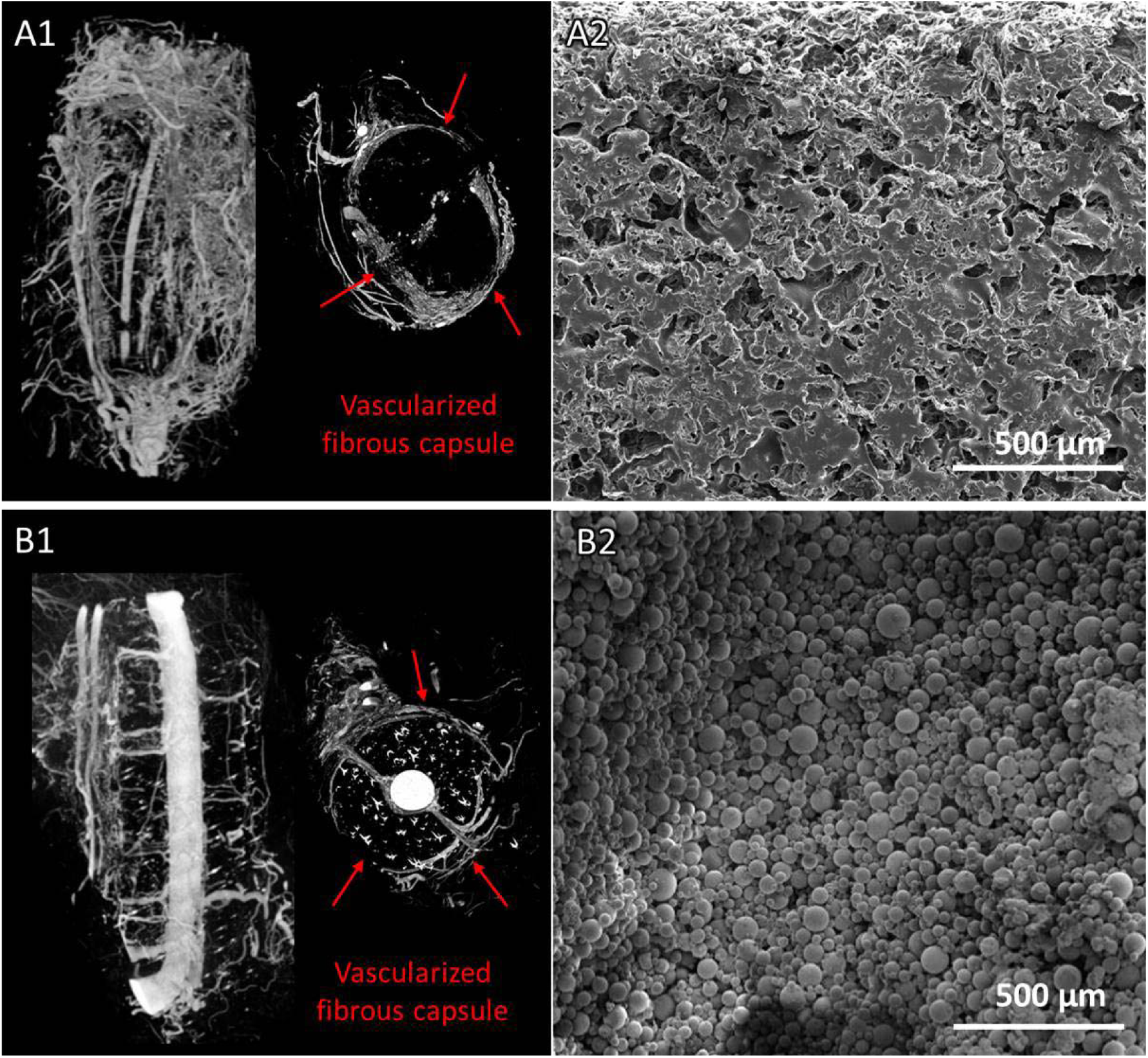
3D micro-CT images of the vascularization generated inside a macroporous PCL scaffold (A1) or a 3D-printed PMMA scaffold (B1), perfused by a vein after 4 weeks of implantation. Most blood vessels are within the fibrous capsule surrounding the implant and not inside the material, as seen in cross section (right). SEM pictures of the surface topography and microporosity of the PCL and PMMA scaffolds (A2 and B2, respectively).

